# Compositae-ParaLoss-1272: Complementary sunflower specific probe-set reduces issues with paralogs in complex systems

**DOI:** 10.1101/2023.07.19.549085

**Authors:** Erika R. Moore-Pollard, Daniel S. Jones, Jennifer R. Mandel

## Abstract

**Premise:** The sunflower family specific probe set, Compositae-1061, has enabled family-wide phylogenomic studies and investigations at lower-taxonomic levels by targeting 1,000+ genes. However, it generally lacks resolution at the genus to species level, especially in groups with complex evolutionary histories including polyploidy and hybridization.

**Methods:** In this study, we developed a new Hyb-Seq probe set, Compositae-ParaLoss-1272, designed to target orthologous loci in Asteraceae family members. We tested its efficiency across the family by simulating target-enrichment sequencing in silico. Additionally, we tested its effectiveness at lower taxonomic levels in genus *Packera* which has a complex evolutionary and taxonomic history. We performed Hyb-Seq with Compositae-ParaLoss-1272 for 19 taxa which were previously studied using the Compositae-1061 probe set. Sequences from both probe sets were used to generate phylogenies, compare topologies, and assess node support.

**Results:** We report that Compositae-ParaLoss-1272 captured loci across all tested Asteraceae members. Additionally, Compositae-ParaLoss-1272 had less gene tree discordance, recovered considerably fewer paralogous sequences, and retained longer loci than Compositae-1061.

**Discussion:** Given the complexity of plant evolutionary histories, assigning orthology for phylogenomic analyses will continue to be challenging. However, we anticipate this new probe set will provide improved resolution and utility for studies at lower-taxonomic levels and complex groups in the sunflower family.

## INTRODUCTION

The sunflower family, also known as the daisy family, Asteraceae, or Compositae, is one of the largest flowering plant families making up roughly 10% of all angiosperms. This large and diverse group has presented many challenges for resolving evolutionary relationships and studying diversifications through time and space. Recent phylogenetic work in the family has employed various methods to reconstruct family-level phylogenies to better understand the evolutionary history and relationships of Asteraceae. For example, Huang et al. (2016) used transcriptome data, Zhang et al. (2021) used a combination of transcriptome and whole-genome sequence data, while Mandel et al. (2019) used Target-Enrichment sequencing (also known as Hyb-Seq) with a custom probe set designed to enrich for conserved gene sequences in Asteraceae (Mandel et al., 2014, 2017). This probe set has become popular among researchers studying members of Asteraceae and has enabled investigations at lower taxonomic levels, especially understudied groups (e.g., Lichter-Marck et al., 2020; Thapa et al., 2020; de Lima Ferreira et al., 2022; Siniscalchi et al., 2019, 2023).

Targeted sequence probe sets have grown in popularity over the last 10 years with sets designed to target loci across large plant groups: bryophytes (i.e., mosses; Liu et al., 2019), pteridophytes (i.e., ferns, Wolf et al., 2018), and angiosperms (i.e., Johnson et al., 2019), as well as for specific plant families (i.e., Asteraceae, Mandel et al., 2014, 2017; Fabaceae, Chapman, 2015; Ochnaceae, Shah et al., 2021; Orchidaceae, Eserman et al., 2021). Typically, low-coverage genome-skim and/or transcriptome data have been used to design probe sets (Straub et al., 2012; Weitemier et al., 2014; Folk et al., 2015; Fonseca and Lohmann, 2020); however, genome-skimming is generally not as effective for designing a probe set for nuclear genes, as low-coverage genome skim data typically enriches for organellar genomes and other high-copy genomic sequences in plants (Stull et al., 2013). These genomic regions are often highly conserved and repetitive and are thus less useful for resolving relationships in some groups. Using transcriptome data offers the potential to sequence and select from thousands of loci, enabling the survey of genomic regions with different rates of molecular evolution.

Several tools have recently become available to design targeted sequence probe sets using transcriptome data more easily, such as OrthoFinder (Emms and Kelly, 2019) and MarkerMiner (Chamala et al., 2015). OrthoFinder is a pipeline that identifies orthogroups and/or orthologs in transcriptomes based on sequence similarities across many species (Emms and Kelly, 2015). In return, the output returns a list of exons usable for probe design. One disadvantage to OrthoFinder, and ultimately the transcriptome-only approach, is that without knowledge of intron-exon topology, probes could overlap boundaries and thus would not be effective at sequence capture (McKain et al., 2018). Alternatively, identification of intron-exon boundaries is straightforward in the MarkerMiner tool, which aligns transcriptome data to reference angiosperm genome sequences and returns intron-masked multiple sequence alignments (Chamala et al., 2015; McKain et al., 2018). The general workflow for MarkerMiner compares user-provided transcriptome sequences against reference genomes with known single-copy orthologous genes (e.g., *Arabidopsis thaliana* (L.) Heynh.), drastically reducing the number of paralogous sequences, or ‘paralogs’, retained for each gene. Probe sets designed using this approach have yielded greater phylogenetic resolution in some groups at the family level (e.g., Cactaceae; Acha and Majure, 2022) and genus/species level (e.g., *Euphorbia* L.; Villaverde et al., 2018; *Zanthoxylum* L., Reichelt et al., 2021). Retaining only single-copy orthologs as a result of MarkerMiner can greatly improve species tree inference as paralogs complicate phylogeny building by causing gene tree heterogeneity. If not accounted for properly, this heterogeneity can lead to misleading phylogeny construction and an incorrect interpretation of species relationships (Smith and Hahn, 2021). Thus, removing or limiting the influence of paralogs can eliminate some of the ambiguity and help with inferring phylogenies more accurately.

In this study, we used 48 transcriptomes to generate a new probe set for sequencing orthologous sequences in Asteraceae utilizing MarkerMiner. Our sampling included 45 Asteraceae taxa and three outgroups from across the order Asterales: Calyceraceae, Campanulaceae, and Goodeniaceae. Though Compositae-1061 has been shown to be efficient at higher-and some lower-taxonomic levels within the family, it generally lacks resolution at the genus to species level. Therefore, we designed this probe set with the aim to provide higher resolution at lower-taxonomic levels and help tackle issues with paralogy, especially among complex groups. To do this, we tested the compatibility and efficiency of this new probe set across the entire family by simulating target-enrichment sequencing in silico in six Compositae members spanning across the family. We then used members of the genus *Packera* Á. Löve & D. Löve as a model system to directly test the efficacy of the probe set by sequencing 16 *Packera* and three outgroup taxa using the Compositae-1061 probe set and this newly designed probe set, named Compositae-ParaLoss-1272. We generated phylogenetic trees, compared their topologies, and assessed node support to determine whether Compositae-ParaLoss-1272 provided greater resolution at the genus/species level compared to Compositae-1061.

## METHODS

### Probe Development

To identify single-copy nuclear loci and select regions for target enrichment probe design, transcriptome data from 48 taxa spanning Asterales (Appendix S1; see Supporting Information with this article) were compiled. Three specimens were collected from the Memphis Botanic Garden live collection, of which we were unable to make an herbarium voucher. All 48 samples were used as input for MarkerMiner v. 1.0 (Chamala et al., 2015) using default settings with both *Arabidopsis thaliana* and *Vitis vinifera* L. as reference genomes. MarkerMiner is an open access, bioinformatic workflow that compares user-provided transcriptomes against reference angiosperm genomes with known single-copy orthologous genes that can be used to design primers or probes for targeted sequencing. Orthologous genes are classified as single copy in the reference genomes if they are present across 17 genomes that were previously annotated as part of a systematic survey on duplication resistant genes (De Smet et al., 2013). The resulting genes were complemented against Angiosperm353 (Johnson et al., 2019) and Compositae-1061 (Mandel et al., 2014, 2017) probe sets to ensure only unique sequences remained.

Exons with lengths ranging from 120 - 1,000bp and a minimum variability of two single nucleotide polymorphisms (SNPs) were selected using a custom python script (https://github.com/ClaudiaPaetzold/MarkerMinerFilter). The resulting 3,853 exonic regions, spanning 1,925 genes around 1,112-85,780bp long (Appendix S2), were further processed by MyBaits at Arbor Biosciences (Ann Arbor, Michigan, USA) to produce a set of 120-mer tiled baits that overlap every 60 bases and share an 80% identity when possible, similar to methods used to develop the MyBaits Compositae-1061 kit (Mandel et al., 2014), hereafter referred to as 1061. Additional filtering steps were implemented as follows: 1) sequence clusters containing five or more taxa that did not target species/lineage specific genes/clusters were retained, 2) clusters containing only the reference sequence data were removed, 3) probes with at least three sequences that covered the alignment were retained, and 4) probes with high similarities (80% or 90%) representing only one or two species were collapsed. Finally, two additional loci were added to the probe design: the MADS-box transcription factor *LEAFY* (*LFY*, Weigel et al. 1992) and the transmembrane pseudokinase *CORYNE* (*CRN*, Müller et al., 2008), two conserved single-copy genes that regulate flower development and meristem size, respectively, in Angiosperms. Gene sequences for *LFY* were identified using the tblastx plugin in Geneious Prime v. 2023.0.4 (https://www.geneious.com) with custom *Bidens ferulifolia* (Jacq.) Sweet (cv. Compact Yellow) leaf transcriptome and *Lactuca sativa* L. genome assembly (v.8) blast databases respectively. The *CRN* gene sequence (AT5G13290) came directly from *Arabidopsis thaliana* using The Arabidopsis Information Resource (TAIR, https://www.arabidopsis.org/).

The resulting MyBaits target enrichment kit contains 60,158 120bp-long, in-solution, biotinylated baits based on target sequence information. The final bait panel, Compositae-ParaLoss-1272, consisted of 13,117 probes and 1,272 loci after filtering.

### Simulating capture sequencing across Compositae

We simulated a target-enrichment sequencing run in silico on six published genomes spanning Asteraceae (Figure 1) using Compositae-ParaLoss-1272, hereafter referred to as 1272, and 1061 in the software CapSim (Cao et al., 2018) to investigate the efficiency of this new probe set for recovering loci across the sunflower family. CapSim is a tool that simulates a sequence run in silico with given a genome sequence and probe set as input. The simulated data can be used for evaluating the performance of the analysis pipeline, as well as the efficiency of the probe design.

**Figure 1.**
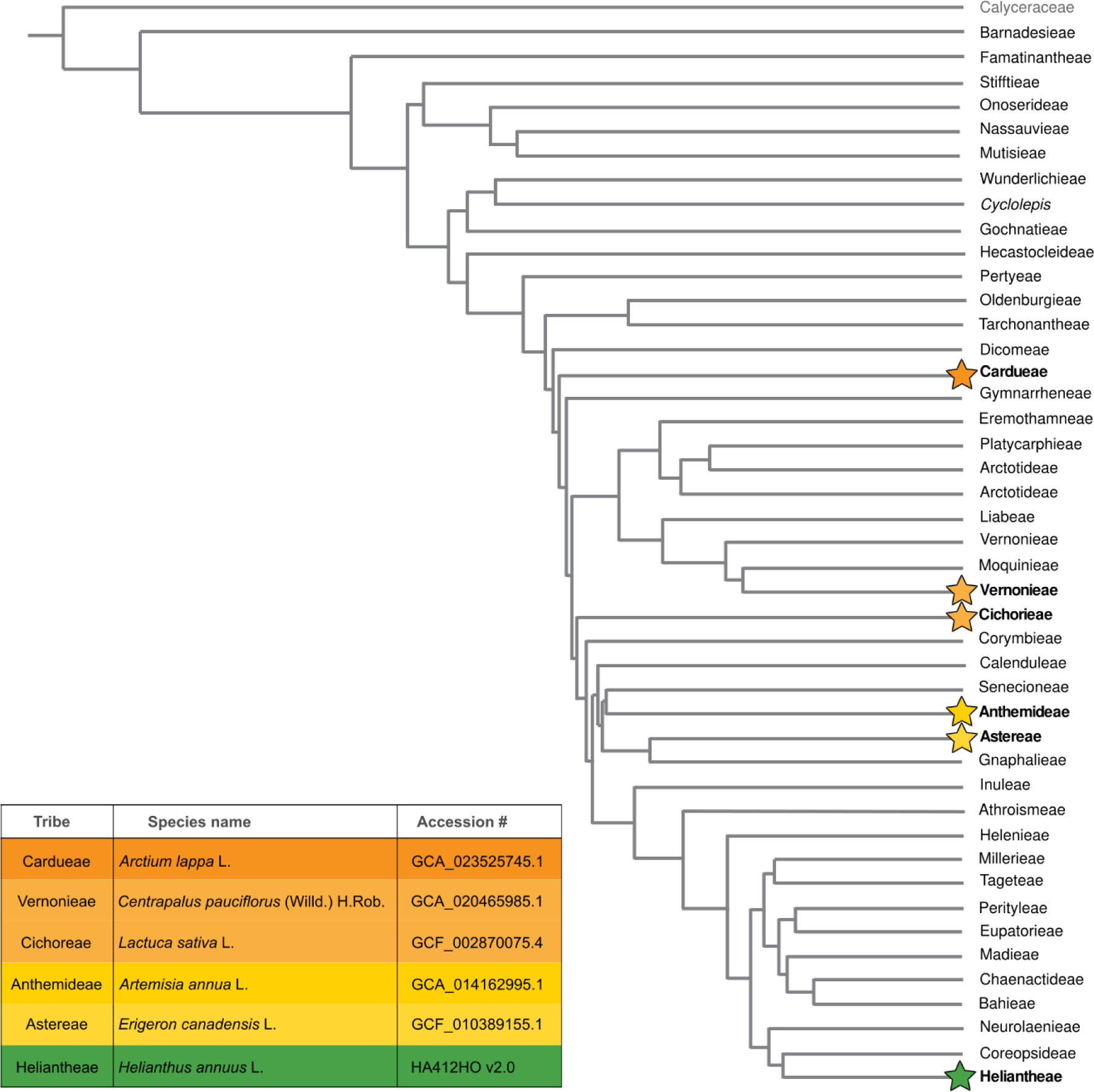
Phylogeny of Asteraceae tribes and the family’s proposed sister group, Calyceraceae, modified from Mandel et al. 2019. Stars at the tip indicate a specimen from that tribe was used for in silico sequencing analyses utilizing CapSim. Colors of stars relate to the table in the bottom left containing sequence accession numbers given by NCBI, excluding *Helianthus annuus* which came from Badouin et al. (2017; https://sunflowergenome.org/assembly-data/).

Prior to running CapSim, an index file was generated, and probes were aligned to the six genomes using Bowtie2 v. 2.3.5.1 (Langmead and Salzberg, 2012; Langmead et al., 2019). After the alignment, the sam files were sorted and indexed into bam files using samtools v. 1.9 (Danecek et al., 2021). The resulting bam files were then used as input in CapSim using the *jsa.sim.capsim* command with the following settings: median fragment size at shearing (-- fmedian) set to 250, miseq simulated (--miseq), illumina read length (--illen) set to 150, and the number of fragments (--num) set to 50,000,000. The resulting fastq files were used as input in the HybPiper v. 2.0.1 (Johnson et al., 2016) pipeline to map simulated sequences against the probe set. Summary and paralog statistics were recovered using the stats and paralog_retriever options in HybPiper.

### Specimen collection

An Illumina sequence run was performed using the new probe set on a selection of 19 total taxa, 16 *Packera* and three outgroup taxa, that were previously sequenced with the 1061 probe set. *Packera* taxa were selected to be representative across the entire *Packera* phylogenetic tree from Moore-Pollard and Mandel (2023). One outgroup taxon, *Packera loratifolia* (Greenm.) W.A.Weber & Á.Löve, was included in this analysis as an outgroup instead of an ingroup since previous studies have shown it is likely misclassified in *Packera* and instead should be in *Senecio* (Barkley, 1985; Bain and Jansen, 1995; Bain and Golden, 2000; Pelser et al., 2007; Moore-Pollard and Mandel, 2023). Illumina sequencing was performed on the 19 taxa utilizing the 1272 probe set following the steps below. A complete list of sampled species, herbarium vouchers, and NCBI accession numbers can be found in Table 1.

**Table 1.** Voucher specimens for the Illumina sequence run. Publication status and authorities assigned by IPNI. * indicates a report for only the Compositae-ParaLoss-1272 probe set.

| Species | Location | Collector and # (Herbarium) | Coll. date | Sheet barcode or ID number | Raw reads (paired)* | Reads mapped* | NCBI accession - 1061 | NCBI accession - 1272 |
| --- | --- | --- | --- | --- | --- | --- | --- | --- |
| *Emilia fosbergii* Nicolson | USA; FL, Osceola County | Wayne D. Longbottom, David H. Williams, Holly L. Williams 14545 (NY) | 18-Nov-10 | 02074297 | 1,572,629,062 | 10,414,762 | SRR22543392 | SRR24860889 |
| *Packera aurea* (L.) Á.Löve & D.Löve | USA; Tennessee, Campbell County | Floden 866 (TENN) | s.d. | N/A | 3,009,238,834 | 19,928,734 | SRR22543326 | SRR24860888 |
| *Packera cana* (Hook.) W.A.Weber & Á.Löve | USA; ID, Adams County | Don Knoke 2101 (WTU) | 25-Jun-11 | 406472 | 4,989,136,942 | 33,040,642 | SRR24862023 | SRR24860878 |
| *Packera candidissima* (Greene) W.A.Weber & Á.Löve | Mexico; Sierra Madre Occidental, Mexico | Robert A. Bye 9680 (ASU) | 26-May-80 | 121438 | 2,880,272,150 | 19,074,650 | SRR22543387 | SRR24860877 |
| *Packera castoreus* (S.L.Welsh) Kartesz | USA; UT, Piute County | Alan Taye 3674 (OSC) | 20-Sep-87 | 172202 | 2,269,567,448 | 15,030,248 | SRR22543385 | SRR24860876 |
| *Packera crocata* (Rydb.) W.A.Weber & Á.Löve | USA; CO, Jackson County | Mary Damm 38 (OSC) | 29-Jul-02 | 244322 | 6,132,282,442 | 40,611,142 | SRR22543379 | SRR24860875 |
| *Packera cynthioides* (Greene) W.A.Weber & Á.Löve | USA; NM, Grant County | Darrell E. Ward 80-010 (NY) | 6-Sep-80 | 03088483 | 2,917,414,224 | 19,320,624 | SRR22543377 | SRR24860874 |
| *Packera dubia* (Spreng.) Trock & Mabb. | USA; NC, Chesapeake County | J. Brandon Fuller (NCU) | 29-Jun-20 | N/A | 2,167,035,730 | 14,351,230 | SRR22543313 | SRR24860880 |
| *Packera franciscana* (Greene) W.A.Weber & Á.Löve | USA; AZ, Coconino County | J. Resinger 1577 (ARIZ) | 14-Jul-76 | 233800 | 4,604,239,452 | 30,491,652 | SRR22543368 | SRR24860873 |
| *Packera glabella* (Poir.) C.Jeffrey | USA; Tennessee, Bradley County | DeSelm 06-04 (TENN) | s.d. | N/A | 3,641,082,026 | 24,113,126 | SRR22543366 | SRR24860872 |
| *Packera greenei* (A.Gray) W.A.Weber & Á.Löve | USA; CA, Trinity County | E.R. Moore 8 (MEM) | 27-Jun-19 | 20904 | 2,943,301,060 | 19,492,060 | SRR22543365 | SRR24860871 |
| *Packera layneae* (Greene) W.A.Weber & Á.Löve | USA; CA, El Dorado County | Kathryn A. Beck 200310 (WTU) | 30-Apr-03 | 375035 | 5,681,052,766 | 37,622,866 | SRR22543356 | SRR24860887 |
| *Packera loratifolia* (Greenm.) W.A.Weber & Á.Löve | Mexico; Sierra La Viga, Mexico | J.A. Villarreal, J. Valdes R 5163 (ASU) | 16-Sep-89 | 182928 | 2,487,875,698 | 16,475,998 | SRR22543355 | SRR24860886 |
| *Packera musiniensis* (S.L.Welsh) Trock | USA; UT, Sanpete County | D. Atwood 21259 (ARIZ) | 9-Aug-96 | 334839 | 2,988,242,284 | 19,789,684 | SRR22543346 | SRR24860885 |
| *Packera porteri* (Greene) C.Jeffrey | USA; OR, County | Coll. Wm. Cusick 2308 (OSC) | 8/3/1899 | 97915 | 4,421,594,684 | 29,282,084 | SRR22543334 | SRR24860884 |
| *Packera pseud aurea* (Rydb.) W.A.Weber & Á.Löve | USA; ID, Valley County | James F. Smith 9147 (OSC) | 29-Jul-10 | 228940 | 3,922,950,102 | 25,979,802 | SRR22543332 | SRR24860883 |
| *Packera streptanthifolia* (Greene) W.A.Weber & Á.Löve | USA; OR, Grant County | Sharon Birks 2010-42 (OSC) | 16-Jul-10 | 255384 | 13,606,754,356 | 90,110,956 | SRR22543319 | SRR24860882 |
| *Packera texensis* O'Kennon<br>& Trock | USA; TX, Gillespie<br>County | B.L. Turner 24-75 (TEX) | 10-Apr-04 | 00211804 | 4,920,515,898 | 32,586,198 | SRR22543316 | SRR24860881 |
| *Roldana gilgii* (Greenm.)<br>H.Rob. & Brettell | Mexico; Chiapas, Mexico | D.E. Breedlove 24411<br>(TEX) | 5-Mar-72 | 00062617 | 2,082,647,568 | 13,792,368 | SRR22543307 | SRR24860879 |

### DNA extraction and sequencing

DNA extraction and sequencing methods followed steps outlined by Moore-Pollard and Mandel (2023). Briefly, dried leaf tissue collected from herbarium specimens was used to extract DNA. DNA length was assessed by running a 1% agarose gel in 1X TBE and GelRed 3x (Biotium), with a target DNA length of 400-500 base pairs (bp). If DNA fragments appeared larger than 500bp, up to 1µg DNA was sheared via sonication with a QSonica machine (ThermoCube, New York, USA). Sheared DNA was then used to generate barcoded libraries utilizing NEBNext Ultra II DNA Library Prep Kit (New England Biolabs, Ipswich, Massachusetts, USA). Libraries produced followed the NEBNext Ultra II Version 5 protocol with size selection on DNA fragments at 300-400bp range but were adjusted by halving the amount of reagents and DNA used to save on supplies and resources. Targeted sequence capture was performed on the libraries using the newly designed probe set, 1272, from Arbor Biosciences (Ann Arbor, Michigan, USA) described above, following manufacturer’s protocols (version 4.01). Captured targets were amplified and quantified using KAPA library quantification kits (Kapa Biosystems, Wilmington, Massachusetts, USA). Quality and quantity checks were performed throughout using a Nanodrop 2000 (Thermo Fisher Scientific, Carlsbad, California, USA) and Qubit High Sensitivity assay (ThermoFisher Scientific, Oregon, USA), respectively. The pooled libraries were sequenced on an Illumina NovaSeq6000 at HudsonAlpha Institute of Technology (Huntsville, Alabama, USA).

### Ortholog assembly and phylogenetic analyses

Raw sequence reads from 1061 and 1272 were cleaned and trimmed of adapters using Trimmomatic v. 0.36 (Bolger et al., 2014), implementing the Sliding Window quality filter (illuminaclip 2:30:10, leading 20, trailing 20, sliding window 5:20). Cleaned reads were retained if they had a minimum length of 36 bp. Cleaned reads were then mapped to the corresponding 1061 (Mandel et al., 2014) or 1272 probe sets using the HybPiper pipeline. A combined reference/*de novo* assembly was performed using BWA v. 0.7.17 (Li and Durbin, 2009) and SPAdes v. 3.5 (Bankevich et al., 2012), respectively, with specified kmer lengths: 21, 33, 55, 77, and 99. Maximum likelihood trees were then built in RAxML v. 8.1.3 (Stamatakis, 2014) with 1,000 bootstrap replicates under the GTR+I+Γ model. Species trees were generated from each resulting RAxML gene matrix using ASTRAL-III v. 5.7.3 (Zhang et al., 2018), a pseudo-coalescent tree building method. Local posterior probability (LPP) values were generated at each node to indicate the probability that the resulting branch is the true branch given the set of input gene trees. LPP is considered a more reliable clade support measure than bootstrapping since it is computed based on a quartet score (Sayyari and Mirarab, 2016) and assumes incomplete lineage sorting (Zhang et al., 2018). The resulting trees, 1061 and 1272, were then visualized using the package *phytools* (Revell, 2012) in R v. 4.0.5 (R Core Team, 2016; RStudio, 2020).

### Measuring phylogenomic discordance

To determine if 1272 increased node resolution across *Packera*, Quartet Sampling (Pease et al., 2018) was used to assess the confidence, consistency, and informativeness of internal tree relationships. Quartet Sampling provides a more comprehensive support value estimate than LPP by calculating four scores, three at each node (quartet concordance [QC], quartet differential [QD], and quartet informativeness [QI]) and one at the tip (quartet fidelity [QF]), to determine if the internal relationships are caused by a lack of data, underlying biological processes, or rogue taxa. QC specifies how often a concordant quartet is inferred over other discordant quartets as a range from -1 to 1: -1 indicates that the quartets are more often discordant than concordant and 1 indicates that all quartets are concordant. QD reveals how skewed the discordant quartets are as a range from 0 (high skew) to 1 (low skew). QI suggests how informative the quartets are as a range from 0 (none are informative) to 1 (all are informative). Each terminal branch is then given a QF score which reports how often a taxon is included in the concordant topology given a range of 0 (taxon is present in none) to 1 (taxon is present in all). Quartet Sampling requires a concatenated nucleotide matrix and a rooted species tree. The concatenated matrices were generated using FASconCAT-G v. 1.02 (Kück and Longo, 2014) into a phylip format. The input phylogeny was then rooted using the *pxrr* command in Phyx (Brown et al., 2017).

PhyParts v. 0.0.1 (Smith et al., 2015) was then used to quantify and visualize discordance in the final phylogenies. PhyParts summarizes and visualizes conflict among gene trees given the resulting species tree topology by performing a bipartition analysis, which helps determine if the node support values are misleading because of underlying discordance. This tool requires a rooted final species tree and rooted gene trees as input. Thus, these trees were rooted to the three outgroup taxa, *Roldana gilgii* (Greenm.) H.Rob. & Brettell, *Emilia fosbergii* Nicolson, and *Packera loratifolia*. The script “phypartspiecharts.py” (available at https://github.com/mossmatters/MJPythonNotebooks) was then used to map pie charts onto the nodes in the final species tree, detailing whether there is one dominant topology in the gene trees with not much conflict, if there is one frequent alternative topology, or many low-frequency topologies.

To estimate similarity scores between the 1061 and 1272 tree topologies, we calculated the adjusted Robinson-Foulds (RF_adj_) distance as outlined by Moore-Pollard and Mandel (2023) between the two trees using the RF.dist function in package *phangorn* (Schliep, 2011) in R. Unrooted ASTRAL-III trees were used as input with the “normalize” argument set to TRUE. RF_adj_ calculates the distance between two unrooted trees, with resulting RF_adj_ values closer to zero indicating that the tree topologies are similar, and values closer to one show complete dissimilarity.

## RESULTS

### CapSim

CapSim results showed that both the 1061 and 1272 probe sets were applicable across a broad range of Asteraceae members since both probe sets retained a moderate number of loci. The 1061 probe set generally retained more loci than 1272 with an average of about 551 loci retained using the 1061 probe set, and an average of 453 loci with the 1272 probe set (Table 2). Even so, the average length of the loci was much longer in the 1272 probe set with genes averaging 1,922bp long, and the 1061 probe set produced genes averaging 403bp long (Appendix S3). Additionally, 1272 produced fewer paralog warnings than 1061 with a range of 0-2 paralogs retained per sample with the 1272 probe set, and a range of 96-250 paralogs per sample with 1061 (Table 2). A full list of statistics can be found in Appendix S3.

**Table 2.** Summary statistics of the CapSim run after running the ‘stats’ function in HybPiper.

| Species | 1061 |  |  |  |  | 1272 |  |  |  |  |
| --- | --- | --- | --- | --- | --- | --- | --- | --- | --- | --- |
|  | Reads mapped | % on target | Genes mapped | % genes retained | Paralog warnings | Reads mapped | % on target | Genes mapped | % Genes retained | Paralog warnings |
| *Artemisia annua* L. | 93,739,367 | 93.7% | 407 | 38.4% | 108 | 97,421,399 | 97.4% | 433 | 34.0% | 1 |
| *Helianthus annuus* L. | 97,351,903 | 97.4% | 750 | 70.7% | 250 | 97,357,378 | 97.4% | 403 | 31.7% | 1 |
| *Centrapalus pauciflorus*<br>(Willd.) H.Rob. | 94,823,613 | 94.8% | 466 | 43.9% | 101 | 97,708,408 | 97.7% | 468 | 36.8% | 0 |
| *Lactuca sativa* L. | 98,218,579 | 98.2% | 749 | 70.6% | 223 | 97,532,753 | 97.5% | 519 | 40.8% | 2 |
| *Erigeron canadensis* L. | 95,987,418 | 96.0% | 548 | 51.6% | 96 | 97,231,893 | 97.2% | 500 | 39.3% | 1 |
| *Arctium lappa* L. | 92,956,716 | 93.0% | 388 | 36.6% | 103 | 97,530,647 | 97.5% | 399 | 31.4% | 1 |

### *Packera* sequence stats

Illumina sequencing resulted in a total of 501 million reads and 76 billion sequences across the 19 newly sequenced taxa utilizing the 1272 probe set. The minimum and maximum number of reads ranged from 10.4 million in *Emilia fosbergii* to 90.1 million in *Packera streptanthifolia* (Greene) W.A.Weber & Á.Löve. (Table 1). The HybPiper pipeline retained 1,049 genes (out of 1,061) when using the 1061 probe set, and 1,213 genes (out of 1,272) with the 1272 probe set. The number of loci recovered for each taxon ranged from 923 in *Packera musiniensis* (S.L.Welsh) Trock to 1,051 in *Roldana gilgii* using the 1061 probe set, and 1,258 in *Packera musiniensis* to 1,271 in *Packera streptanthifolia* using the 1272 probe set. The number of loci retained was proportionally higher in 1272 compared to 1061 (Figure 2B), though the 1061 alignment contained fewer missing data (1061: 34.89%; 1272: 35.05%) and was more parsimony informative (1061: 11.7%; 1272: 8.3%) than 1272 (Appendix S4). Alternatively, the 1272 probe set recovered drastically fewer paralogous sequences (‘paralogs’) than the 1061 probe set, with only about 5% of the recovered loci reporting as paralogous, compared to 59% with the 1061 probe set (Figure 2A). The number of paralog warnings ranged from 35-407 genes per sample with the 1061 probe set, compared to 0-14 in the 1272 probe set (Table 3). Additionally, 1272 recovered much longer loci compared to 1061 (Figure 3). Refer to Appendix S4 for a full compilation of statistics.

**Figure 2.**
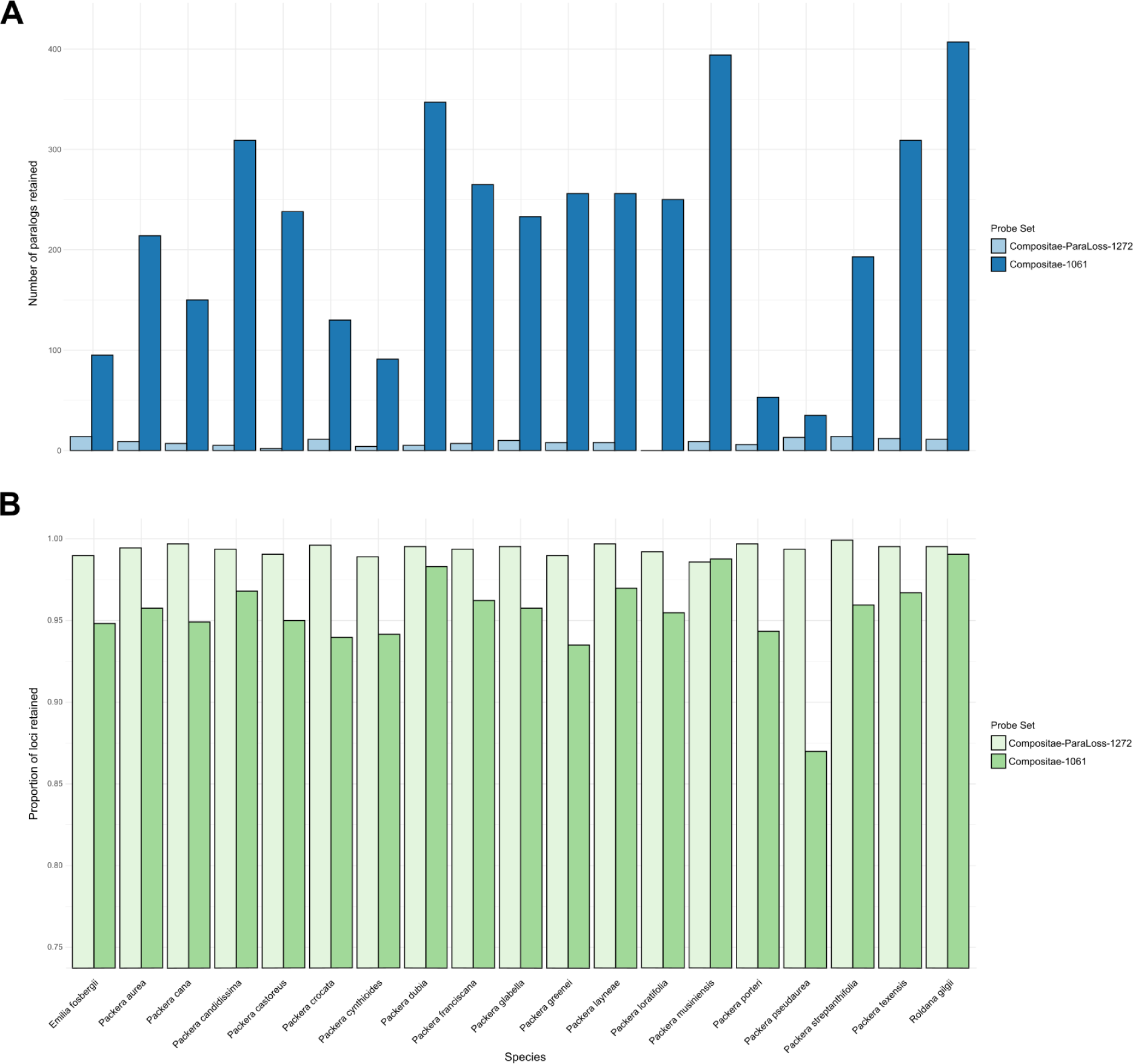
Barplots showing the **A**) number of flagged paralogs, and **B**) the proportion of loci retained for each species dependent on the probe set used. Lighter colors represent the Compositae-ParaLoss-1272 probe set, while darker colors represent the Compositae-1061 probe set as indicated by the keys to the right of the plots. Barplots were generated using base R.

**Figure 3.**
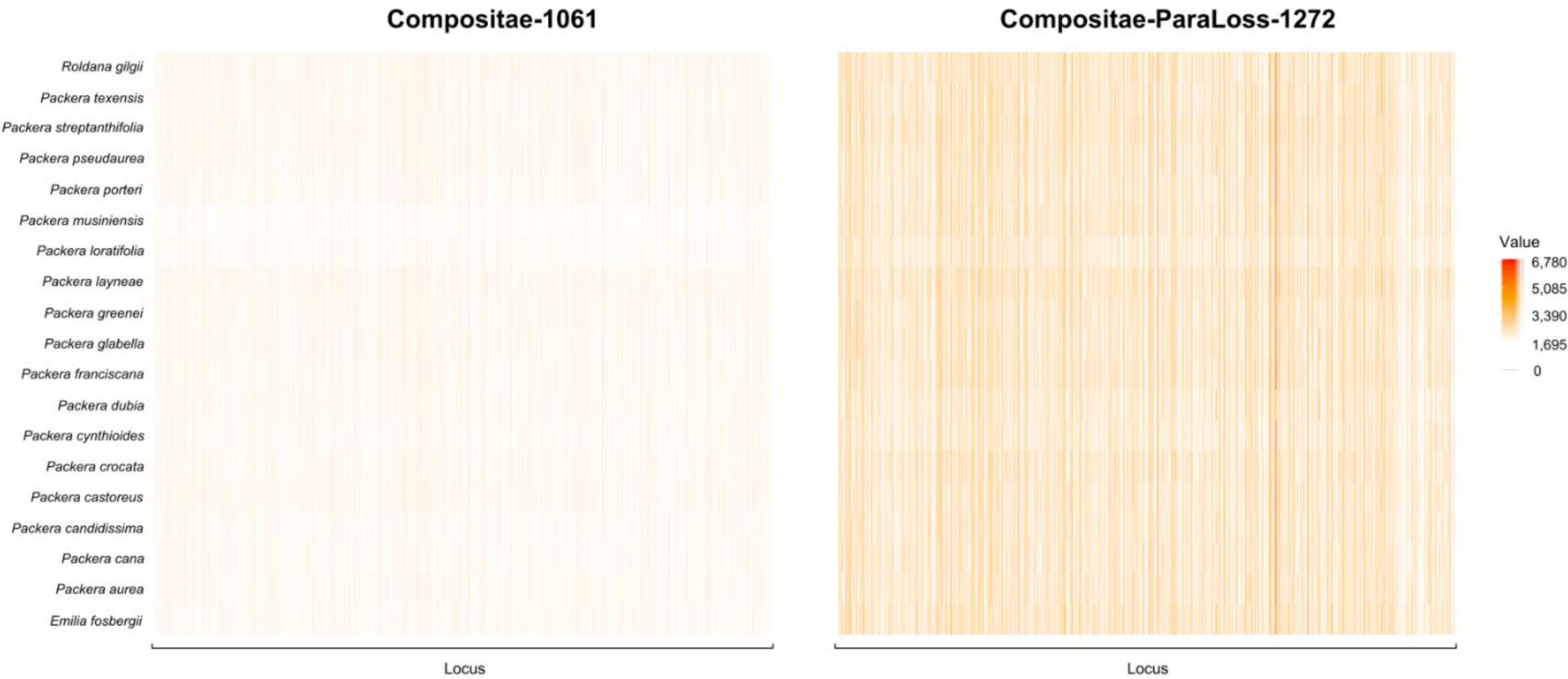
Heatmap of retained locus length in the Compositae-1061 (left) and Compositae-ParaLoss-1272 (right) analyses for each locus (x-axis) of every species (y-axis). The longest loci are indicated by vertical red lines with the smallest loci indicated by vertical orange lines. Loci not retained are shown as white. Heatmaps were generated in R.

**Table 3.** Summary statistics of the Illumina sequencing run after running the ‘stats’ function in HybPiper.

| Species | 1061 |  |  |  |  | 1272 |  |  |  |  |
| --- | --- | --- | --- | --- | --- | --- | --- | --- | --- | --- |
|  | Reads mapped | % on target | Genes mapped | % genes retained | Paralog warnings | Reads mapped | % on target | Genes mapped | % genes retained | Paralog warnings |
| *Emilia fosbergii* Nicolson | 1,185,704 | 59% | 1006 | 94.8% | 95 | 11,236,129 | 65% | 1259 | 99.0% | 14 |
| *Packera aurea* (L.) Á.Löve & D.Löve | 5,184,671 | 53% | 1016 | 95.8% | 214 | 4,438,388 | 26% | 1265 | 99.4% | 9 |
| *Packera cana* (Hook.) W.A.Weber & Á.Löve | 1,532,039 | 21% | 997 | 94.0% | 130 | 8,297,634 | 35% | 1268 | 99.7% | 7 |
| *Packera candidissima* (Greene) W.A.Weber & Á.Löve | 558,742 | 36% | 999 | 94.2% | 91 | 5,690,438 | 43% | 1264 | 99.4% | 5 |
| *Packera castoreus* (S.L.Welsh) Kartesz | 1,654,718 | 37% | 1043 | 98.3% | 347 | 2,912,785 | 32% | 1260 | 99.1% | 2 |
| *Packera crocata* (Rydb.) W.A.Weber & Á.Löve | 2,361,884 | 36% | 1021 | 96.2% | 265 | 10,762,526 | 35% | 1267 | 99.6% | 11 |
| *Packera cynthioides* (Greene) W.A.Weber & Á.Löve | 1,171,793 | 29% | 1007 | 94.9% | 150 | 2,064,556 | 36% | 1258 | 98.9% | 4 |
| *Packera dubia* (Spreng.) Trock & Mabb. | 4,514,739 | 39% | 1016 | 95.8% | 233 | 2,775,445 | 26% | 1266 | 99.5% | 5 |
| *Packera franciscana* (Greene) W.A.Weber & Á.Löve | 1,573,692 | 41% | 992 | 93.5% | 256 | 9,169,648 | 45% | 1264 | 99.4% | 7 |
| *Packera glabella* (Poir.) C.Jeffrey | 1,972,057 | 34% | 1029 | 97.0% | 256 | 6,012,371 | 31% | 1266 | 99.5% | 10 |
| *Packera greenei* (A.Gray) W.A.Weber & Á.Löve | 2,024,706 | 34% | 1013 | 95.5% | 250 | 4,102,840 | 27% | 1259 | 99.0% | 8 |
| *Packera layneae* (Greene) W.A.Weber & Á.Löve | 2,814,096 | 35% | 1048 | 98.8% | 394 | 8,240,509 | 26% | 1268 | 99.7% | 8 |
| *Packera loratifolia* (Greenm.) W.A.Weber & Á.Löve | 511,859 | 43% | 1001 | 94.3% | 53 | 2,435,806 | 35% | 1262 | 99.2% | 0 |
| *Packera musiniensis* (S.L.Welsh) Trock | 68,064 | 9% | 923 | 87.0% | 35 | 6,518,518 | 44% | 1254 | 98.6% | 9 |
| *Packera porteri* (Greene) C.Jeffrey | 1,510,836 | 39% | 1018 | 95.9% | 193 | 5,896,137 | 40% | 1268 | 99.7% | 6 |
| *Packera pseud aurea* (Rydb.) W.A.Weber & Á.Löve | 3,914,039 | 41% | 1027 | 96.8% | 309 | 7,423,481 | 41% | 1264 | 99.4% | 13 |
| *Packera streptanthifolia* (Greene)<br>W.A.Weber & Á.Löve | 2,695,188 | 38% | 1026 | 96.7% | 309 | 23,885,890 | 39% | 1271 | 99.9% | 14 |
| *Packera texensis* O'Kennon &<br>Trock | 2,516,755 | 33% | 1008 | 95.0% | 238 | 9,026,502 | 38% | 1266 | 99.5% | 12 |
| *Roldana gilgii* (Greenm.) H.Rob.<br>& Brettell | 1,545,552 | 28% | 1051 | 99.1% | 407 | 2,105,459 | 34% | 1266 | 99.5% | 11 |

### Discordance of *Packera* taxa

A higher number of gene trees were represented in the final 1272 species tree compared to the 1061 tree (Normalized quartet score = 0.461 and 0.424, respectively). Additionally, the 1272 probe set provided higher resolution at internal nodes compared to the previous probe set, with 13 of the 17 internal nodes having local posterior probability (LPP) values greater than or equal to 0.97LPP, eight of those being fully supported (1.0LPP). This is compared to the 1061 probe set which only had eight nodes greater than or equal to 0.97LPP, seven of those with 1.0LPP (Figure 4). Additionally, the level of discordance of internal *Packera* relationships varied between both trees. Quartets are more often discordant than concordant in the 1061 tree, with four internal nodes having negative Quartet Concordance (QC) values, compared to only one node (between *Packera pseudaurea* (Rydb.) W.A.Weber & Á.Löve and *P. aurea* (L.) Á.Löve & D.Löve, QC = -0.3) in the 1272 tree (Figure 5).

**Figure 4.**
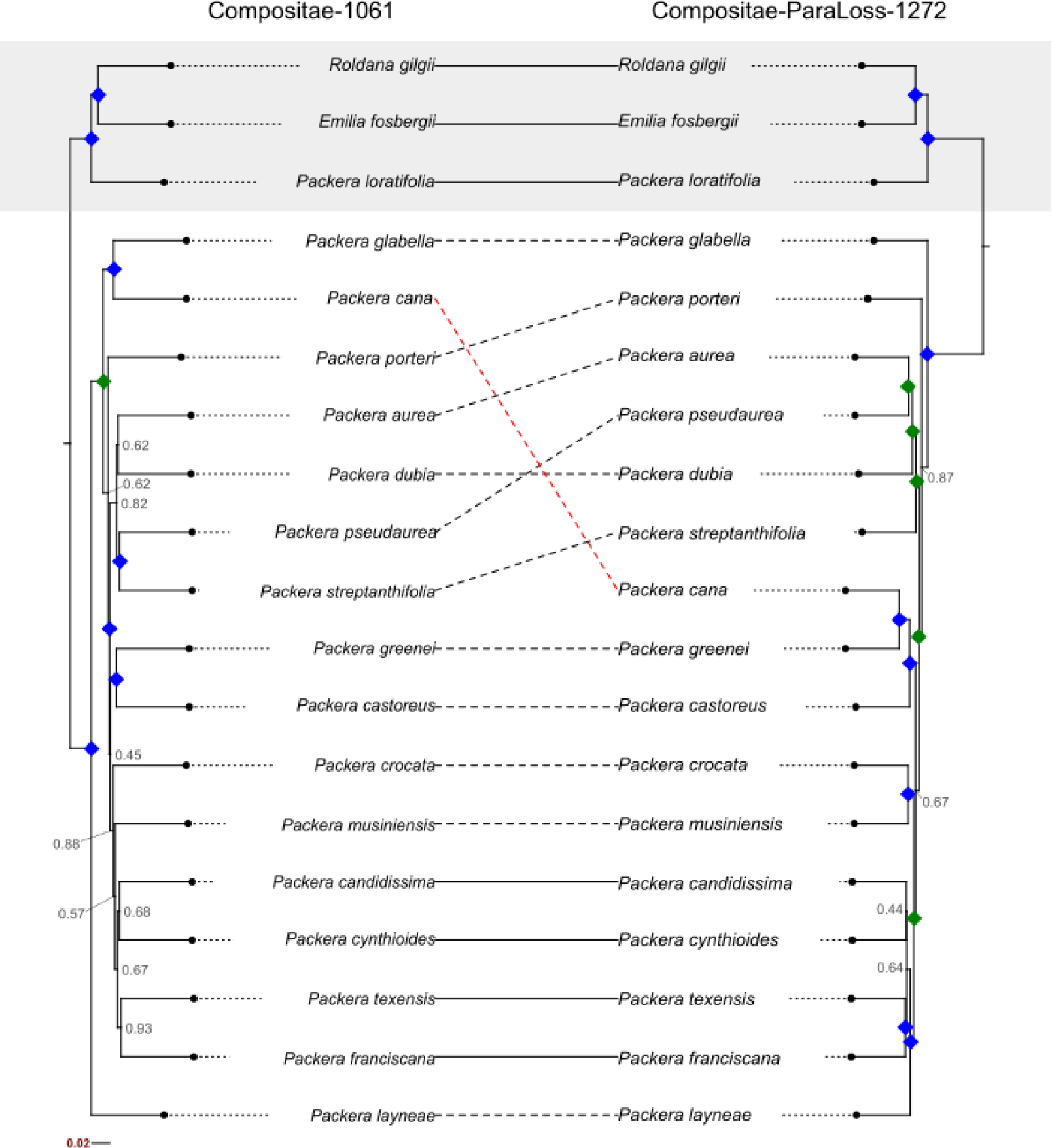
Tanglegram comparing species topologies when phylogenies were developed using the Compositae-1061 probe set (left) or the Compositae-ParaLoss-1272 probe set (right). Completely concordant topologies are indicated with a solid line, medium discordance is indicated by a dashed line, and major discordance is indicated by a dashed, red line. Local posterior probability (LPP) values of 1.0LPP are indicated by a blue diamond at the node. LPP values ranging from 0.97-0.99 are indicated by a green diamond. All other LPP values lower than 0.97 are shown at the corresponding node. Outgroup species are highlighted with a gray shadow box.

**Figure 5.**
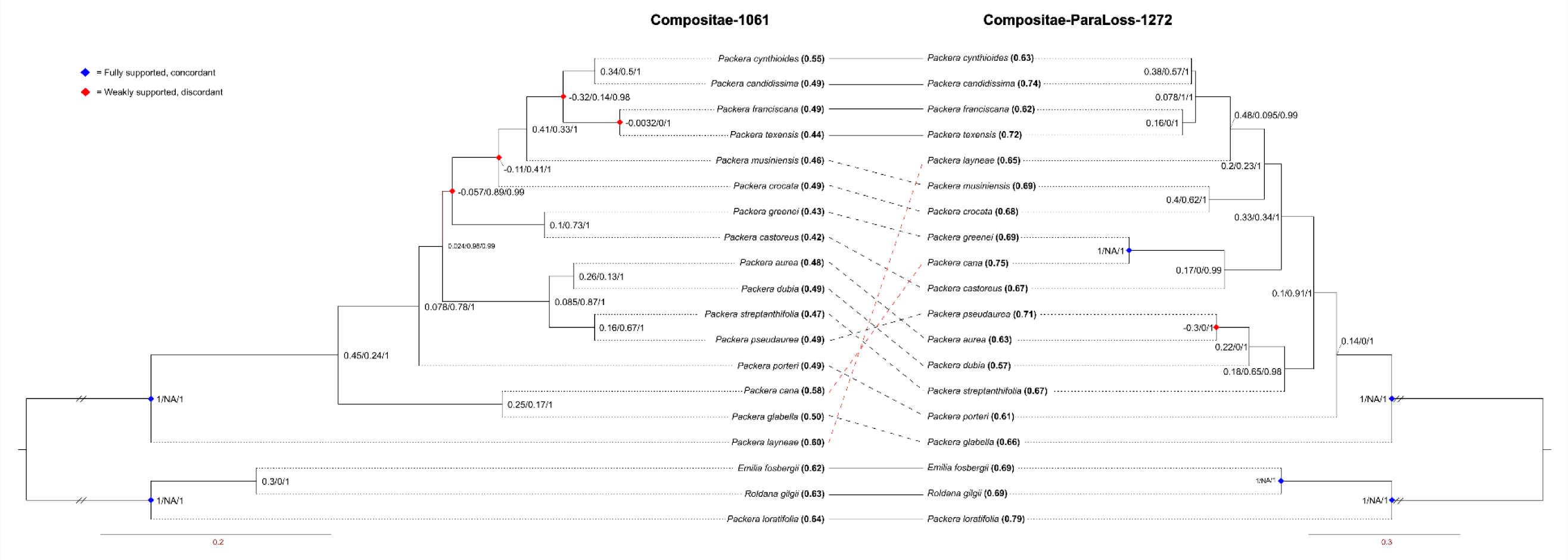
Discordance and support values in the Compositae-1061 (left) and Compositae-ParaLoss-1272 (right) trees indicated by Quartet Sampling. At each node, three values are represented: Quartet Concordance (QC), Quartet Differential (QD), and Quartet Informativeness (QI), shown as QC/QD/QI. Blue diamonds at node indicate fully supported and concordant quartets, red diamonds indicate weakly supported and discordant quartets. Quartet Fidelity (QF) scores are at each tip label in parenthesis and bolded.

The resulting 1061 and 1272 species tree topologies were moderately incongruent with each other (RF_adj_ = 0.625). Of the taxon relationships that remained the same in both trees, 1272 showed more concordant and strongly supported relationships compared to 1061 (Figures 5 and 6). For example, both tree topologies have *P. cynthioides* (Greene) W.A.Weber & Á.Löve and *P. candidissima* (Greene) W.A.Weber & Á.Löve as sister, and *P. franciscana* (Greene) W.A.Weber & Á.Löve and *P. texensis* O’Kennon & Trock as sister; all four within the same smaller clade (Figure 5). However, the node between *P. franciscana* and *P. texensis* and the node joining the two sister groups were majorly discordant in the 1061 tree (QC = -0.0032, -0.32; respectively), while the same relationships in the 1272 tree were less discordant (QC = 0.16, 0.078; respectively). Even so, the relationships were still not strongly supported. The outgroup relationships and monophyly of *Packera* were fully supported in the 1272 tree (Figure 5). Alternatively, the 1061 tree showed the monophyly of *Packera* with full support; however, the relationship between the outgroup taxa, *Emilia fosbergii* and *Roldana gilgii,* showed weak support with a discordant skew (QS score at node: 0.3/0/1; Figure 5). Quartet fidelity (QF) scores were generally higher in the 1272 tree than the 1061 tree, which ranged from 0.57-0.79 and 0.42- 0.64, respectively (Figure 5), indicating a higher percentage of quartet topologies involving the tested taxa were concordant with the focal tree branch in the 1272 tree.

**Figure 6.**
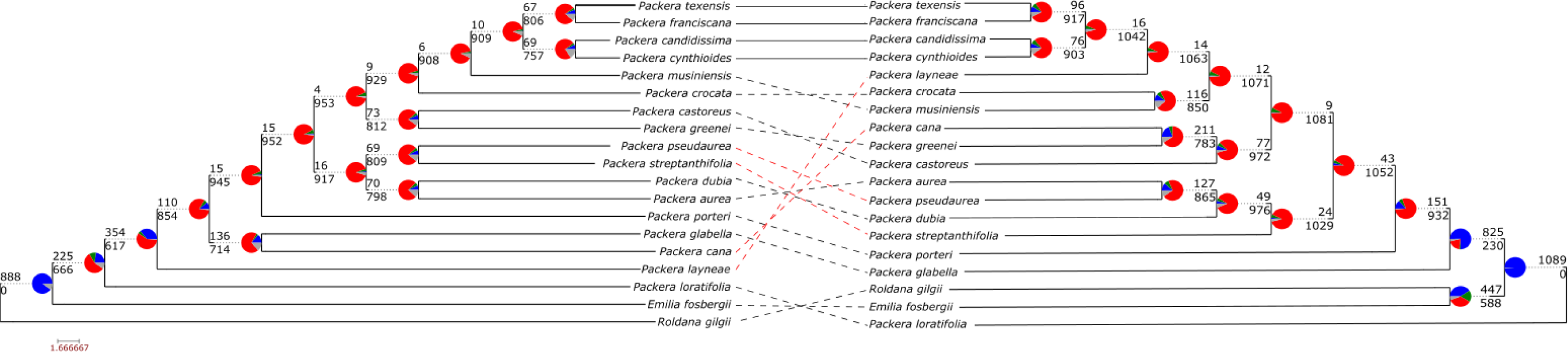
PhyParts results between the Compositae-1061 probe set (left) and Compositae-ParaLoss-1272 probe set (right). Pie charts at nodes show the percentage of gene tree discordance or concordance when compared to the final species tree. The color scheme reveals the percentage of gene trees that are: concordant (blue), the top alternative bipartition (green), all other alternative bipartitions (red), or uninformative at that node (gray). Numbers above and below the branch indicate the number of concordant (blue) and conflicting (red) gene trees, respectively.

## DISCUSSION

In this study, we designed and tested a complimentary Compositae-specific probe set, Compositae-ParaLoss-1272, that provided higher resolution at the lower-taxonomic levels of species in our *Packera* test case. The new probe set dramatically reduced the number of paralogs recovered, retained longer gene sequences, and was likely important for improving the resolution in our *Packera* comparison. Our new probe set was also applicable across all tested members of Asteraceae and recovered more and longer orthologous genes than 1061 (Appendix S3), as well as retained a substantially lower number of paralogs than 1061 (Table 3) when tested in silico. Additionally, there is the ability to do a double capture since the genes associated with 1061 and Angiosperm-353 are not included in the 1272 probe design.

When comparing the 1061 and 1272 tree topologies to the larger *Packera* phylogeny (Moore-Pollard and Mandel, 2023), the 1272 tree’s evolutionary relationships was in higher agreement with the whole-genus phylogeny (RF_adj_ = 0.6) as compared to 1061 (RF_adj_ = 0.667), potentially indicating this new probe set is more robust to species sampling compared to 1061. For example, our 1061 tree places *P. layneae* (Greene) W.A.Weber & Á.Löve as sister to the remaining core *Packera* species. This relationship differs from both the 1272 and Moore-Pollard and Mandel (2023) trees, which have *P. layneae* placed more internally and with other Californian endemic species (Figure 4; Moore-Pollard and Mandel, 2023). Additionally, the placement of *P. glabella* (Poir.) C.Jeffrey in the 1061 tree differs from past phylogenomic studies, including the 1272 tree in this study, which place it as sister to all remaining *Packera* taxa (Freeman, 1985; Barkley, 1988; Trock, 1999; Bain and Golden, 2000; Schilling and Floden, 2015). While this is promising, further studies are needed to investigate whether the new probe set is more robust to taxon sampling.

The resulting tree topologies between 1061 and 1272 were moderately incongruent (RF_adj_ = 0.625; Figure 4), indicating that species relationships varied dependent on the probe set used. We suggest that these differences can be explained by 1) the different gene sets used to make the phylogeny, 2) the differences in paralog retention, or 3) underlying biological processes. First, given that this new probe set was complemented against Compositae-1061 during production, there is no overlap of gene sequences between probe sets so only unique gene sequences, which have their own evolutionary histories, were used to generate each phylogeny. Therefore, the tree topologies and species relationships could differ since the 1272 phylogeny may be reflecting unique gene histories not shared with 1061, and vice versa. Next, having fewer paralogs, as is seen in 1272, removes ambiguity of the data so the resulting species relationships in the 1272 tree may better reflect “true” relationships as there is not as much underlying gene heterogeneity (Smith and Hahn, 2021; Zhou et al., 2021). Finally, underlying biological processes, such as hybridization, reticulation, or incomplete-lineage sorting (ILS), may be influencing our results as these processes are known to cause complications in phylogenetic construction (Arnold, 1997; Maddison, 1997; Alberts et al., 2002; Nussbaum et al., 2007).

Although only marginal, the 1272 tree had lower levels of discordance, indicating that 1272 provides better resolution at the nodes than 1061 (Figures 5 and 6), though the nodes are still highly discordant. It is reasonable to consider that these underlying biological processes may be influencing our concordance results as *Packera* members have a long history of reticulation (e.g., Bremer, 1994; Bain et al., 1997) and hybridizing in the wild (e.g., Fernald, 1943; Barkley, 1962; Chapman et al., 1971; Uttal, 1984; Bain, 1988; Trock, 1999; Gramling, 2006; Weakley et al., 2011). Similar conclusions have also been found in other groups (e.g., Sessa et al., 2012; Vargas et al., 2017; Morales-Briones et al., 2018). Another possible explanation for the low node resolution is that only a subset of taxa (16 out of 88 *Packera* taxa) were used to generate these phylogenies. Having such low species sampling could influence species relationships and node support values given a lack of data (Heath et al., 2008; Sanderson et al., 2010).

Alternatively, 1272 contained more missing data and was considered less parsimony informative (PI) than 1061 (Appendix S4); however, the differences were minimal (PI_1272_ = 23.4%, PI_1061_ = 24.1%). Interestingly, similar results were found in a previous study that generated a Fabaceae specific probe set using MarkerMiner and compared the results to other probe design methods (Vatanparast et al., 2018). This study found that MarkerMiner produced fewer paralogous loci than other design methods, but also was not as parsimony informative as other methods, following our results.

While our results showed that 1061 retained a higher number of genes in silico (Table 2), the Illumina sequencing run of the 1272 probe set shows much higher locus retention and greater resolution than the 1061 probe set (Table 3). We hypothesize that the low loci retention in silico is a relic of read simulators not always capturing the variances of Illumina sequenced data since they cannot model noise or sequencing technology biases perfectly (May et al., 2022; Duncavage et al., 2023). Additionally, we suspect that having longer gene sequences in the probe set influences read simulator results, though we cannot confirm the validity of these suspicions.

Ultimately, the most notable difference between the 1272 and 1061 probe sets is the number of paralogs retained per individual, which was far fewer in the 1272 probe set than the 1061. Given this, the 1061 and 1272 probe sets are still comparable options for target-enrichment sequencing in lower-taxonomic members of Compositae. However, the low paralog retention of the 1272 probe set can be very advantageous when dealing with groups known to be complicated by polyploidy since polyploidy is typically associated with higher paralog retention (Lynch and Conery, 2000; Wolfe, 2001; Veitia, 2005). More attention is being placed on polyploidy in non-model plant groups (e.g., Lim et al., 2008; Bellinger et al., 2022; Fernández et al., 2022), and the underlying issues associated with it are becoming more well known (see Rothfels, 2021). Being able to address these issues early in the phylogenomic pipeline can improve phylogenetic reconstructions and provide more confidence in data interpretations. Given this, we anticipate that further testing will provide additional support for the utility of this probe set in complex groups in the sunflower family. We hope this design approach will be seen as a model for other complex systems.

## ACKNOWLEDGEMENTS

The authors thank Matthew D. Pollard for his bioinformatic help. We also thank Brian Brunelle at Arbor Biosciences for his help and expertise with the probe design. Additionally, we thank the University of Memphis High-Performance Cluster (HPC) administrators, Eric Spangler and Kristian Skjervold, for their assistance with the HPC and willingness to help. Finally, we thank Jane Grimwood at HudsonAlpha.

## AUTHOR CONTRIBUTIONS

E.R.M.P. designed the probe set, generated and analyzed data, and wrote the manuscript. J.R.M. helped design the probe set. D.S.J. provided transcriptome data for probe design and funds for sequencing. J.R.M. and D.S.J. provided edits to the manuscript.

All authors approved of the final version.

## DATA AVAILABILITY STATEMENT

Raw sequence data are available in the National Center for Biotechnology Information (NCBI) Sequence Read Archive (Bioprojects: PRJNA978591 and PRJNA994483).

Additional Supporting Information may be found online in the Supporting Information section at the end of the article.

**Appendix S1.** Voucher specimens to develop the probe set using MarkerMiner. Species names and authorities assigned by IPNI.

**Appendix S2.** List of 1,925 targeted loci in the Compositae-ParaLoss-1272 probe set and information about their associated functions in *Arabidopsis thaliana* L. (source: The *Arabidopsis* Information Resource (TAIR); https://www.arabidopsis.org/tools/bulk/genes/index.jsp). *Vitis vinifera* L. specific genes that have no known function (n = 17) are included.

**Appendix S3.** Table of HybPiper summary statistics for the six Asteraceae genomes from the CapSim run.

**Appendix S4.** General and full HybPiper stats of the Illumina sequence run.

**Appendix S5.** Compositae-ParaLoss-1272 probe set file for bioinformatic analyses.

## Notes

### Competing Interest Statement

The authors have declared no competing interest.

